# Reversal of splicing infidelity is a pre-activation step in B cell differentiation

**DOI:** 10.1101/2022.05.13.491723

**Authors:** Tina M. O’Grady, Melody Baddoo, Samuel A. Flemington, Nate A. Ungerleider, Erik K. Flemington

## Abstract

B cell activation and differentiation is central to the adaptive immune response. Changes in exon usage can have major impacts on cellular signaling and differentiation but have not been systematically explored in differentiating B cells. We analyzed exon usage and intron retention in RNA-Seq data from subsets of human B cells at various stages of differentiation, and in an *in vitro* laboratory model of B cell activation and differentiation (Epstein Barr virus infection). Blood naïve B cells were found to have an unusual splicing profile, with unannotated splicing events in over 30% of expressed genes. Splicing changed substantially upon naïve B cell entry into secondary lymphoid tissue and before activation, involving significant increases in exon commitment and reductions in intron retention. These changes preferentially involved short introns with weak splice sites and were likely mediated by an overall increase in splicing efficiency induced by the lymphoid environment. The majority of transcripts affected by splicing changes showed restoration of encoded conserved protein domains and/or reduced targeting to the nonsense-mediated decay pathway. Affected genes were enriched in functionally important immune cell activation pathways such as antigen-mediated signaling, cell cycle control and mRNA processing and splicing. Functional observations from donor B cell subsets in progressive states of differentiation and from timecourse experiments using the *in vitro* model suggest that these widespread changes in mRNA splicing play a role in preparing naïve B cells for the decisive step of antigen-mediated activation and differentiation.

## Introduction

B-cell recognition of antigens and the production of antibodies lie at the core of the adaptive immune response. B cells originate in the bone marrow via a finely tuned pathway that optimizes breadth and specificity of antigen-binding capacity while avoiding autoreactivity. Upon maturation, antigen-naïve B cells (NBCs) with functional B cell receptors (BCRs) emerge from the bone marrow into the bloodstream, and eventually migrate towards secondary lymphoid tissues such as tonsils, adenoids and lymph nodes. The secondary lymphoid tissues are immune system hubs where the initially nonproliferative NBCs may encounter their antigens and cognate T cells in a signaling environment that promotes activation. Upon antigen binding to the BCR and T cell stimulation, B cells initiate an elaborate cascade of signaling events. This leads to a rapid and orchestrated increase in the transcription and translation of genes required to induce clonal expansion and the formation of a germinal center (GC). In the GC, proliferating B cells undergo somatic hypermutation to produce BCRs with increased antigen affinity, and class-switch recombination to generate mature antibody subtypes. Successfully mutated and class-switched GC B cells undergo further transcriptomic and phenotypic changes and differentiate into long-lived antibody-secreting plasma cells (PCs).

Alternative splicing of mRNA has been found to be linked to many cellular signaling and differentiation processes, with highly specific splicing profiles associated with different tissue types and cell subsets in the immune system and elsewhere (1). Differential abundance or activity of splicing factors can lead to skipping or inclusion of exons, retention of introns, or usage of alternate splice sites. Splicing differences, in particular intron retention variations, have been reported in subsets of neural (2), muscle (3), hematologic (4-6) and immune (7-11) cells.

We used RNA-Seq datasets from human donors to thoroughly interrogate splicing in B cell subsets across a spectrum of differentiation, including NBCs in venous blood, NBCs in secondary lymphoid tissue, GC B cells and fully differentiated PCs. We identified splicing changes over the course of differentiation that have extraordinary penetrance in the B cell transcriptome, affecting over 30% of expressed genes. Early in the differentiation pathway, blood-derived NBCs have a unique splicing profile, marked by high intron retention and a large number of aberrant exon skipping events that are corrected through the course of differentiation. Surprisingly, most splicing changes occur upon NBC entry into the secondary lymphoid tissue, before antigen-mediated activation induces proliferation and GC formation. These early, pre-proliferative splicing changes are similarly observed in a controlled laboratory model of human B cell activation (*in vitro* Epstein Barr virus (EBV) infection of blood NBCs), further supporting a broad increase in splicing fidelity as a prelude to B cell activation.

## Results

### Pervasive unannotated splicing in blood naïve B cells

To investigate RNA splicing in human B cell subsets, we obtained high read-count, paired-end RNA-Seq data from fluorescence activated cell-sorted (FACS) human donor cells via the Blueprint Epigenome project ((12) **Supplemental file 1**). Assessing splicing across these datasets, we noticed a remarkably large number of unusual junctions that were not annotated in the Ensembl reference transcriptome, particularly in datasets derived from blood NBCs. To investigate these novel splicing events in more detail, we used stringent criteria to select high confidence unannotated splice junctions. For each robustly expressed gene (transcripts per million (TPM) > 10), we first identified the annotated splice junction with the highest read count. Only unannotated splicing events with a read count of at least 10% of the depth of the gene’s highest annotated splice junction were further considered (**Supplemental file 2**). Using these criteria, we found that over 30% of genes in blood NBCs from each of the three donors contained at least one unannotated splice junction, noticeably higher than the proportion in lymphoid (tonsil)-derived NBCs, GC B cells, PCs, or the extensively used EBV-immortalized B cell lymphoblastoid cell line (LCL) GM12878 (**Figure 1A**, (13, 14)).

**Figure 1.**
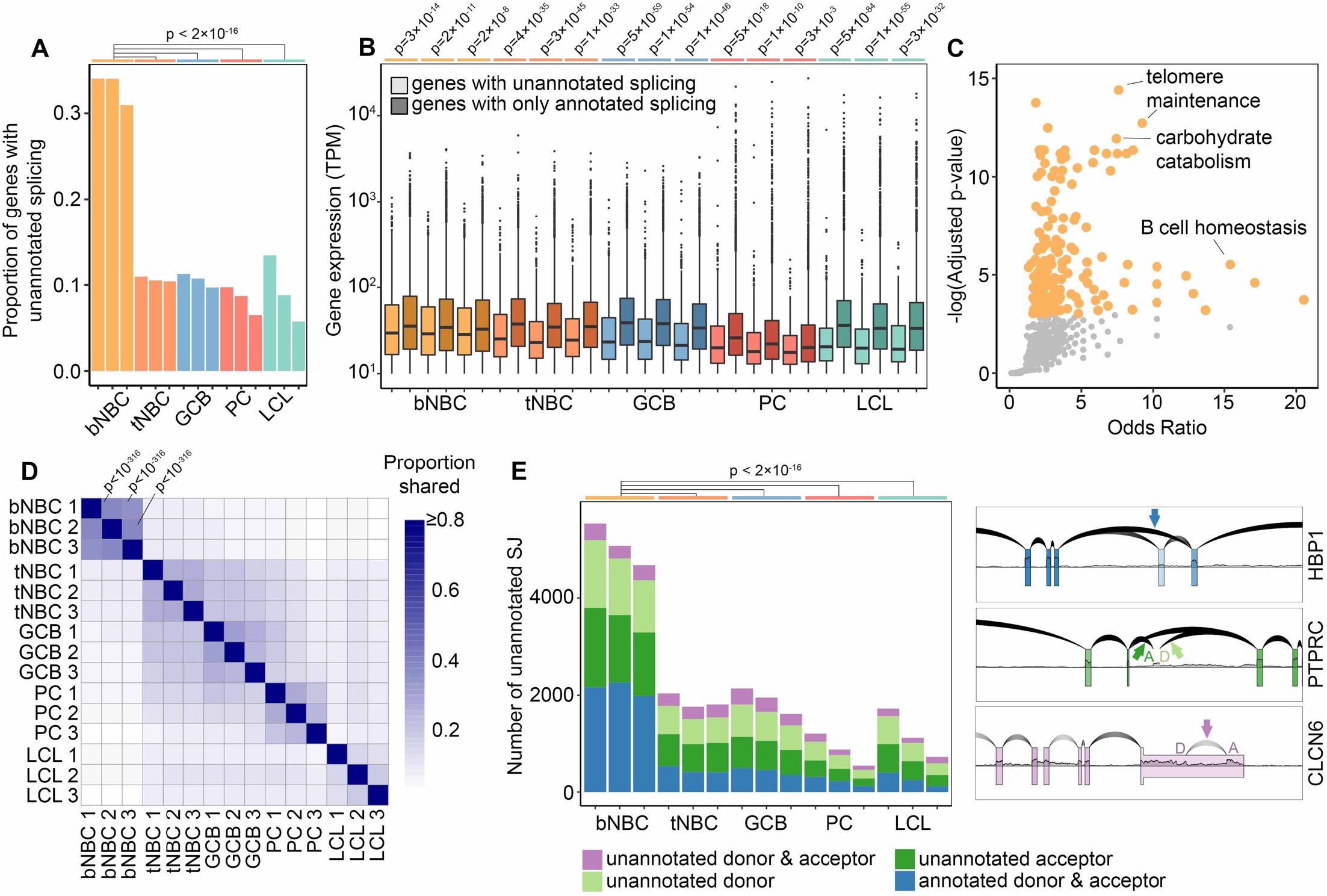
Pervasive unannotated splicing in blood naïve B cells. (A) Proportion of robustly expressed genes (TPM > 10) with high-confidence unannotated splicing in B cell subsets and LCL samples. P-values calculated using pairwise comparison of proportions test with Bonferroni correction. (B) Gene expression level in TPM (minimum = 10) by splice junction annotation status and sample. Lighter colors represent genes containing unannotated splice junctions, darker colors represent genes with annotated splicing only. P-values calculated by Wilcoxon rank sum test with continuity correction and Bonferroni correction. (C) GO Biological Process term enrichment in genes with unannotated splicing in all three blood NBC samples. Orange points = significantly enriched terms (adjusted p-value < 0.05). Grey points = terms not significantly enriched. Adjusted p-values calculated by Enrichr using Fisher’s exact test with Benjamini-Hochberg correction. The complete list of GO terms and adjusted p-values is available as Supplemental file 3. (D) Proportion of shared unannotated splice junctions in pairwise comparisons of all samples. For comparisons between blood NBCs, p-values calculated by hypergeometric test are indicated. Complete p-values are available as Supplemental file 4. (E) Number of unannotated splice junctions using annotated donors, acceptors, both, or neither in each sample of B cells and LCLs (left) and examples of different types of unannotated junctions (right) using RNA-Seq data from blood NBCs plotted with SpliceV. bNBC = blood naïve B cells, tNBC = lymphoid (tonsil) naïve B cells, GCB = germinal center B cells, PC = plasma cells, LCL = lymphoblastoid cell line GM12878.

Genes with unannotated splice junctions were on average expressed at slightly lower levels than genes with entirely annotated junctions, but nevertheless included very highly expressed genes (**Figure 1B**). Notably, however, the most highly expressed genes typically did not contain unannotated splice junctions (**Figure 1B**). This may indicate an essential requirement for specific isoforms of these genes and commensurately strong *cis* splicing features, and/or it may reflect more comprehensive annotation of alternate isoforms in highly expressed genes.

To evaluate functional implications of aberrant splicing, we identified genes in each subset with unannotated splicing across all replicates and examined these gene sets using Enrichr (15). Blood NBC genes with unannotated splicing were strongly enriched in several tissue-specific Gene Ontology (GO) terms such as *GO:0001782: B cell homeostasis*, possibly indicating the presence of novel stage-specific isoforms of mRNA and protein. GO terms such as *GO:0043470: regulation of carbohydrate catabolic process, GO:0043467: regulation of generation of precursor metabolites and energy, GO:0010389: regulation of G2/M transition of mitotic cell cycle* and *GO:0032210: regulation of telomere maintenance via telomerase* were also enriched, potentially representing a limited need for canonical forms of gene products involved in energy production and the cell cycle in these resting, unactivated cells. For the remaining B cell subsets, no term or pathway showed statistically significant enrichment (**Figure 1C, Supplemental file 3**).

To assess whether the extensive unannotated splicing in blood NBCs is a programmed regulatory feature of B cell differentiation, we examined the consistency of unannotated splicing events across multiple donors and B cell subsets. We performed pairwise comparisons of high-confidence unannotated splice junctions in all samples and calculated the proportion of unannotated junctions shared between the samples (**Figure 1D, Supplemental file 4**). Blood NBCs had broad similarity among the three donors and formed a distinct group, separate from the other samples. This high level of consistency indicated that the unique splicing pattern in blood NBCs is an innate, regulated feature of this B cell subtype.

Assessing individual unannotated splicing events in blood NBCs, we noted that many novel splicing events represented exon skipping of one or more constitutive exon(s), with both the splice donor and acceptor sites being annotated (**Figure 1E**, HBP1 example). Also prevalent were hemi-annotated events in which either the donor or the acceptor sites were unannotated (**Figure 1E**, PTPRC example). Lastly, a small percentage of unannotated junctions were completely novel with both the donor and acceptor sites unannotated (**Figure 1E**, CLCN6 example). The varied nature of unannotated splicing events is consistent with compromised activity of multiple splicing factors in blood NBCs, with the skipping observed in HBP1 (**Figure 1E**) potentially arising from diminished U2 complex activity at the acceptor site and/or reduced binding of splicing enhancers to the skipped exon, and splicing into and out of intronic sequences (PTPRC – **Figure 1E**) possibly arising from decreased intronic splicing silencer factor activity. Overall, the diversity of unannotated splicing and the robust numbers of unannotated splicing events in blood NBCs (greater than 4000) are consistent with the hypothesis that splicing fidelity is depressed in blood NBCs due to a diverse restriction of splicing factor function.

### Splicing changes are early events in B cell differentiation

Our finding of an unusually high level of novel splicing in blood NBCs compared to other B cell subsets suggests that this splicing pattern is largely reversed early in differentiation, upon NBC entry into lymphoid tissue. To corroborate this hypothesis, we used the *in vitro* EBV infection model, which mimics activation, proliferation and differentiation of B cells in a controlled tissue culture environment (16, 17). Our analysis of the fully EBV-immortalized LCL GM12878 indicated that unannotated splicing in this cell line was comparable to that of activated, differentiated B cells *in vivo* (**Figure 1**). To investigate the stage at which the unique splicing pattern of blood NBCs is reversed, we obtained RNA-Seq datasets from timecourse experiments of EBV infection leading to LCL establishment (17-20). Consistent with previously observed similarities between *in vitro* EBV-associated activation and *in vivo* antigen-mediated activation (16, 17), fully EBV-immortalized LCLs from two different laboratories showed strong enrichment in the expression of genes associated with PCs relative to their parental blood B cells (**Figure 2A**). Assessing the EBV infection timecourses, we found substantial decreases in unannotated splicing after infection (**Figure 2B**). Strikingly, the splicing changes occurred predominantly by 2 days post-infection (dpi), early in the timecourses (**Figure 2B**, Blood A and Blood B) and well before the previously reported onset of infection-associated cell cycle entry and proliferation (21). Notably, NBCs harvested in another laboratory from lymphoid tissue (adenoids) had less unannotated splicing than blood B cells, consistent with our observations of tonsil NBCs (**Figure 1**) and there were only modest further decreases upon EBV infection (**Figure 2B**, Lymphoid). Infection of a differentiated EBV-negative Burkitt lymphoma (BL) cell line, Akata-EBV-N, had little impact on the already low level of unannotated splicing in these cells (**Figure 2B**, BL). Together, these *in vitro* B cell activation experiments support the findings from our analyses of *in vivo* B cell subtypes and show that reversal of the unusual blood NBC splicing program is an early event, occurring before NBCs enter the cell cycle and likely upon their entry into secondary lymphoid tissue.

**Figure 2.**
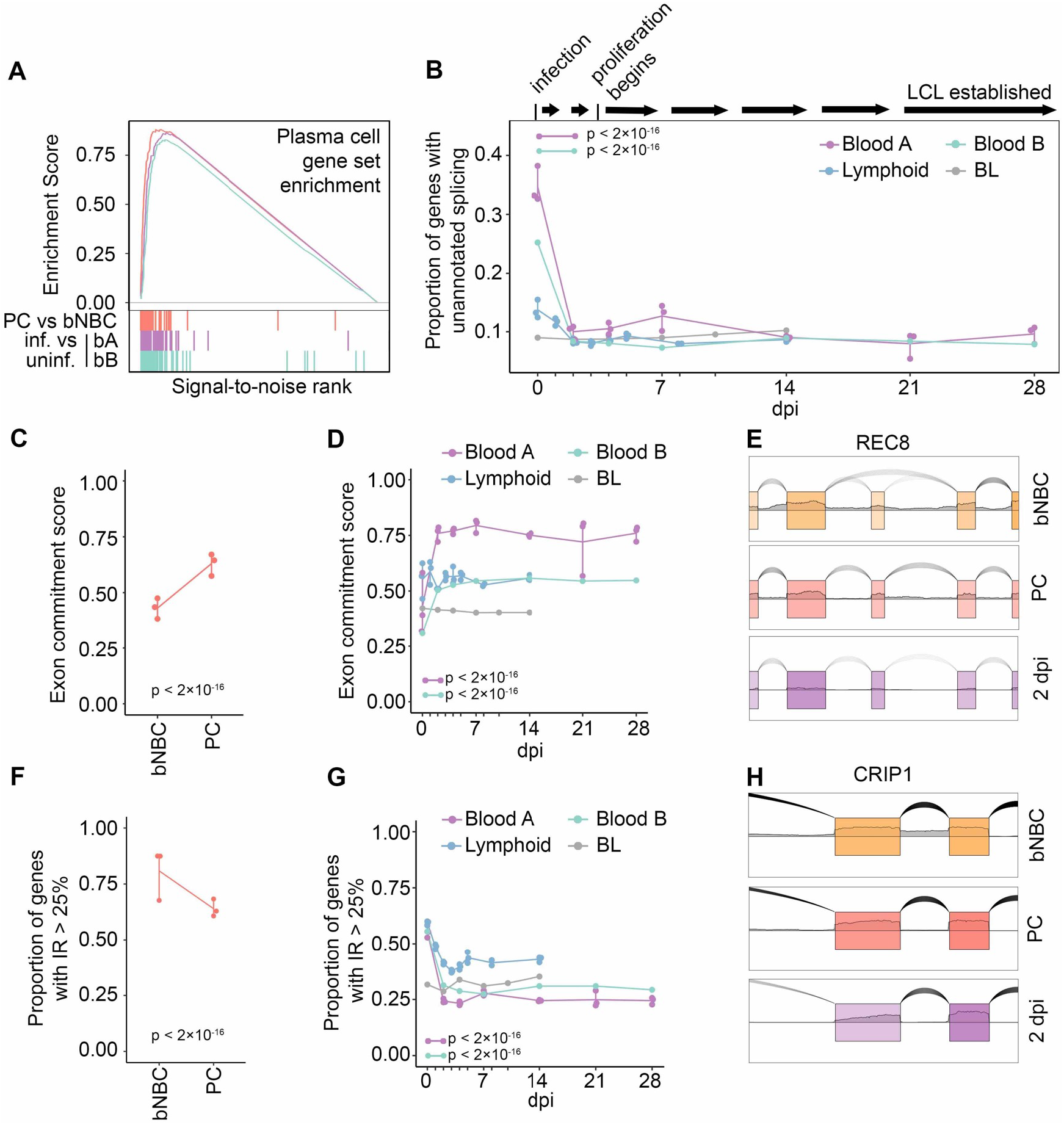
*In vivo* or *in vitro* activation of B cells restores the splicing program. (A) Enrichment scores from GSEA of gene expression level changes in PCs compared to blood NBCs (PC vs bNBC, red, maximum enrichment score = 0.880, p = 0), EBV-infected blood B cells from timecourse A at 28 dpi compared to uninfected cells (bA: purple, maximum enrichment score = 0.863, p = 0) and EBV-infected blood B cells from timecourse B at 28 dpi compared to uninfected cells (bB: green, maximum enrichment score = 0.829, p = 0). Gene set: TARTE_PLASMA_CELL_VS_B_LYMPHOCYTE_UP from MSigDB. Colored vertical lines indicate genes in the gene set. (B) Proportion of genes (TPM > 10) with unannotated splicing after EBV infection. A timeline of EBV infection events is at top. P-values calculated using a pairwise comparison of proportions test with Bonferroni correction are indicated for Blood A and Blood B, 2dpi compared to 0 dpi. (C) Exon commitment score for blood NBC (bNBC) and PC samples. Exon commitment score = proportion of exons either fully included (rMATS IncLevel > 0.99) or fully excluded (rMATS IncLevel < 0.01) in genes with TPM > 10. P-value calculated using 2-sample test for equality of proportions with continuity correction. (D) Exon commitment scores after EBV infection. P-values calculated using a pairwise comparison of proportions test with Bonferroni correction are indicated for Blood A and Blood B timecourses, 2dpi compared to 0 dpi. (E) Example of uncommitted exon in blood NBCs that is committed by increased inclusion upon differentiation to PC or EBV infection (2 dpi, Blood A), plotted with SpliceV. (F) Proportion of genes containing introns with high retention (IRFinder IRratio > 0.25) in blood NBC (bNBC) and PC samples. P-value calculated using 2-sample test for equality of proportions with continuity correction.… (G) Proportion of genes containing introns with high retention levels for at multiple timepoints after EBV infection. P-values calculated using a pairwise comparison of proportions test with Bonferroni correction are indicated for Blood A and Blood B, 2dpi compared to 0 dpi. (H) Example of intron retention in blood NBCs reduced upon differentiation to PC or EBV infection (2 dpi, Blood A), plotted with SpliceV. Blood A = blood infection timecourse A, 3 replicates, Blood B = blood infection timecourse B, 1 replicate, Lymphoid = adenoid infection timecourse, 3 replicates. BL = Akata-EBV-N Burkitt lymphoma cell infection timecourse, 1 replicate.

### Low exon commitment and high intron retention in blood naïve B cells

In general, unannotated splicing in blood NBCs was robust but not fully penetrant, i.e. a portion of transcripts from each affected gene was aberrantly spliced with the remaining transcripts displaying conventional splicing (**Figure 1E, Figure S1**). To globally investigate alternative splicing penetrance, we used rMATS to assess the inclusion level of exons across the transcriptome. For each sample we calculated an “exon commitment” score, which we define as the proportion of exons that were either completely spliced-in (i.e. included in >99% of transcripts from the gene) or completely spliced out (i.e., included in <1% of transcripts from the gene). In blood NBCs exon commitment scores were low, illustrating a lack of commitment to specific isoforms. Upon differentiation exon commitment increased, with a majority of exons in PCs being committed to either inclusion or exclusion (**Figure 2C, S2A**). In the EBV infection timecourses, uninfected blood B cells showed even lower exon commitment, which was substantially reversed by EBV infection. In contrast, no change in exon commitment was observed after infection of lymphoid NBCs or differentiated BL cells (**Figure 2D, S2B-D**). While exon commitment scoring takes into consideration both commitment to the spliced-in and the spliced-out isoforms, it is notable that for most exons, commitment entailed a strong bias toward splicing-in upon differentiation (**Figure 2E, Figure S2A-D**).

NBCs have previously been reported to have high intron retention in both humans and mice (7). RNA-Seq datasets from the Blueprint Epigenome project are not poly(A)-selected and include nascent RNAs with incomplete splicing, making discerning true intron retention events in mature mRNA less precise. Nevertheless, we observed the previously reported decreases in intron retention in differentiated cells relative to blood NBCs in these datasets (**Figure 2F, S3A, (7)**). The poly(A)-selected RNA-Seq data from the EBV infection timecourse experiments, which reflect only mature RNA transcripts, showed even more striking decreases in intron retention after infection of blood B cells (**Figure 2G-H, S3B-D**). The changes in lymphoid NBCs after EBV infection were substantially less than those in blood B cells, and BL cells did not show appreciable intron retention changes upon infection. Overall, the aberrant splicing program in blood

NBCs is marked by low exon commitment and high intron retention, affecting a broad swath of genes. While canonical transcripts are produced for most expressed genes, the extensive splicing infidelity, in the form of mature transcripts with retained introns or skipped exons, likely dampens output of functional proteins until NBCs encounter the lymphoid environment and prepare for activation.

### Involvement of splicing regulatory factors in differentiating B cells

To investigate the basis for the observed splicing infidelity in blood NBCs, we compared differences in exon skipping events between blood NBCs and lymphoid NBCs, GC B cells and PCs to changes in exon skipping observed after knockdown of each of 182 RNA-binding proteins (RBPs), including known and potential splicing regulators (22). RNA-seq datasets for each pair of control or RBP-siRNA transfections in the lymphoblast cell line K562 were downloaded from ENCODE (23, 24) and statistically significant differences in exon skipping were analyzed using rMATS. Strikingly, substantially greater numbers of differential exon skipping events were observed when comparing blood NBCs to lymphoid NBCs, GC B cells, PCs or EBV-infected cells than after the knockdown of any splicing factor tested, including the core acceptor factors U2AF1 and U2AF2 (**Figure S4A**). We then performed pairwise comparisons of altered splicing events in each RBP knockdown experiment to altered splicing events in blood NBCs vs each B cell subset and in uninfected blood B cells vs each EBV infection timepoint. Knockdown of several RBPs led to statistically significant overlap in differential exon skipping events with blood NBCs as compared to lymphoid NBCs, PCs, or EBV-infected cells. These included U2AF1, U2AF2, and other U2 acceptor factors (e.g. SF3B4 and SF3B1), the splicing enhancer SRSF1, and splicing suppressors such as HNRNPC and HNRNPA2B1 (**Figure 3A and Supplemental file 5**). The greatest overlap was observed with knockdown of AQR, a component of the spliceosome that is involved in small nucleolar ribonucleoprotein (snoRNP) assembly. As a negative control, we compared overlap in changed exon skipping events after artificially reversing the direction of change for blood NBCs vs lymphoid NBCs, GC B cells, and PCs and observed minimal overlap with changes detected in RBP knockdown experiments (**Figure S4B**). Our findings that the knockdown of any tested splicing factor is insufficient to recapitulate the number or type of changes in exon skipping upon blood NBC differentiation is consistent with a coordinated set of RBP changes leading to a broad, general increase in splicing fidelity as an early step in B cell differentiation.

**Figure 3.**
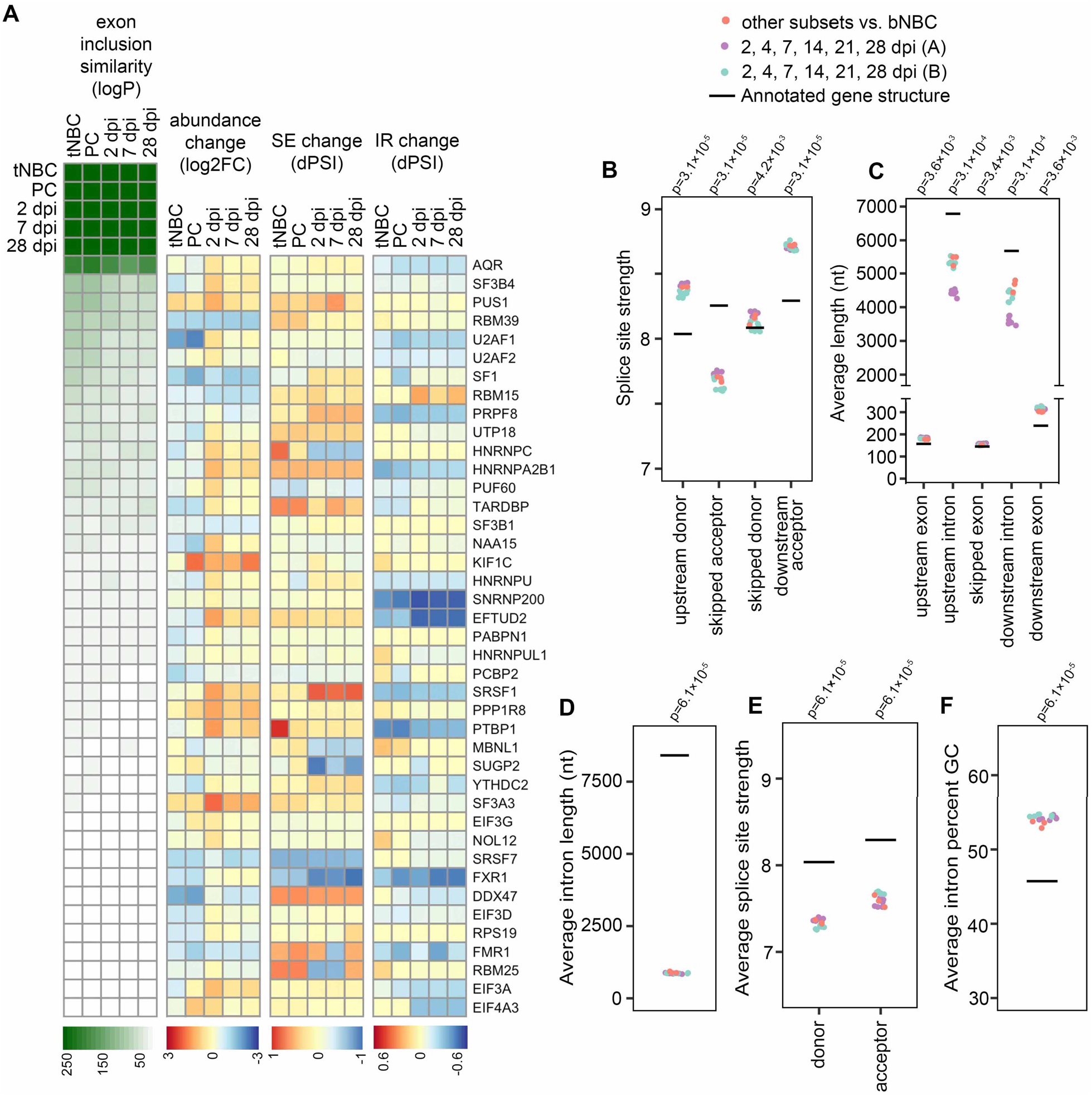
Increased splicing program efficiency alters the splicing profile. (A) Heatmaps indicating exon inclusion similarity (logP = -log(p-value) by hypergeometric test), mRNA abundance change (log2FC = log_2_(fold change) by sleuth), skipped exon (SE) change (dPSI = difference in percent spliced-in by rMATS), and intron retention (IR) change (dPSI by IRFinder) for RBPs with exon inclusion changes most similar to those of B cell differentiation/EBV infection. For RBP genes with multiple splice junctions the SE and IR events with the highest absolute dPSI value measurable in all compared replicates are displayed. PC = PCs compared to blood NBCs; tNBC = lymphoid (tonsil) NBCs compared to blood NBCs; 2, 7 and 28 dpi = EBV-infected cells at specified timepoints compared to uninfected cells in Blood timecourse A. (B) Average splice site strength calculated by MaxEntScan for exons showing increased inclusion in differentiating or EBV-infected B cells (FDR < 0.05 by rMATS). (C) Average length in nucleotides (nt) of exons showing increased inclusion in differentiating or EBV-infected B cells (FDR < 0.05 by rMATS), and their flanking introns and exons. P-values compared to annotated averages calculated by Wilcoxon signed rank exact test with Bonferroni correction. (D) Average length in nucleotides (nt) of introns showing decreased retention in differentiating or EBV-infected B cells (FDR < 0.05 by rMATS). … (E) Average splice site strength calculated by MaxEntScan for introns showing decreased retention in differentiating or EBV-infected B cells (FDR < by rMATS). (F) Average GC content of introns showing decreased retention in differentiating or EBV-infected B cells (FDR < 0.05 by rMATS). Blood infection timecourse A (purple) includes 6 timepoint comparisons with 3 replicates each, Blood infection timecourse B (green) includes 6 timepoint comparisons with 1 replicate each, B cell subset comparisons include 3 subsets (tonsil NBCs, GC B cells, and PCs) compared to blood NBCs. Each B cell subset includes 3 replicates. For panels B-F, p-values compared to annotated averages were calculated by Wilcoxon signed rank exact test with Bonferroni correction.

Notably, some RBPs showing functional relationships with exon skipping differences in blood NBCs vs differentiating cells showed modest increases in mRNA abundance during differentiation (**Figure 3A**). Others showed changes in exon inclusion or decreases in intron retention that likely lead to more efficient production of functional splicing factors (**Figure 3A and Supplemental file 5**). A parallel analysis of overlap in intron retention events in RBP knockdowns and blood NBCs vs differentiating cells also implicated a group of possible RBP effectors of intron retention changes, with many of these showing increased abundance, less exon skipping or less intron retention (**Figure S4C and Supplemental files 5 & 6**). Altogether, the splicing changes upon RBP knockdowns combined with the abundance and splicing changes of RBPs themselves is consistent with a general increase in splicing efficiency occurring alongside the known increase in transcription (25) as B cells progress from resting, unactivated cells to activated, proliferating and/or antibody-secreting cells.

### Weaker splice sites and shorter introns are associated with missplicing in blood naïve B cells

Exon-skipping junctions specific to blood NBCs most often skipped a single exon, though some junctions spanned multiple exons (**Figure S4D**). The Maximum Entropy Model (26) splice site strengths for skipped exons had a distinctive pattern, with an unusually weak acceptor site for the skipped exon flanked by unusually strong upstream exon donor and downstream exon acceptor sites (**Figure 3B**). This is consistent with limited availability of active acceptor recognition factors in blood NBCs causing reduced spliceosome engagement with the weak skipped acceptor but minimal impacts at the strong upstream donor and downstream acceptor sites. Interestingly, blood NBC skipped exons also tended to reside between introns that were relatively short (**Figure 3C**), a characteristic that has been linked to suboptimal transcription rates (27), a property previously reported in naïve T cells (28) that may also be a feature of blood NBCs.

Introns with decreased retention in differentiating B cells also had distinct characteristics. They were substantially shorter than average, with weaker donor and acceptor splice sites, and higher GC content (**Figure 3D-F**). These features are all known to be associated with intron retention both in basal conditions and during splicing impairment (29-31), and have previously been observed in retained introns in mouse NBCs (7). Overall, these observations suggest an inefficient splicing program in blood NBCs with decreased recognition of sub-optimal splice sites.

### B cell exon commitment restores conserved protein domains and evades nonsense-mediated mRNA decay

Alternative splicing events can have drastically different downstream effects in different contexts. Skipped coding exons with lengths divisible by 3 (in-frame skipped exons) remove sequence without disrupting the reading frame. The resulting protein isoforms may serve distinct functions in the cell, may be nonfunctional, or may even act as dominant negative isoforms if a regulatory domain is spliced out. Out-of-frame skipped exons disrupt open reading frames, potentially introducing a premature stop codon (PTC) and targeting the transcript for degradation via the nonsense-mediated decay (NMD) pathway (32).

We investigated the impact of exon skipping events by first determining if the skipped exon was in-frame or out-of-frame. For out-of-frame exons, we scanned the new reading frame downstream of the alternatively spliced exon for PTCs. Transcripts with PTCs more than 50 nt upstream of splice junctions were identified as likely NMD targets, according to previously identified NMD-targeting characteristics (32). A large proportion of skipped exons in blood NBCs appeared to target their transcripts to NMD, while a very small proportion of skipped exons in more differentiated cells were NMD-targeting (**Figure 4A**). For in-frame skipped exons, we determined their encoded amino acid sequences and investigated whether they included conserved protein domains (CD) using the NCBI’s Batch CD-Search tool (33). The proportion of skipped exons in blood NBCs that encoded a conserved domain was much higher than that in differentiated cells. The high number of NMD-targeting and domain-deleted transcripts in blood NBCs suggests that the early splicing program that produces nonfunctional, potentially deleterious transcripts is repaired during B cell differentiation, leading to restored inclusion of critical exons.

**Figure 4.**
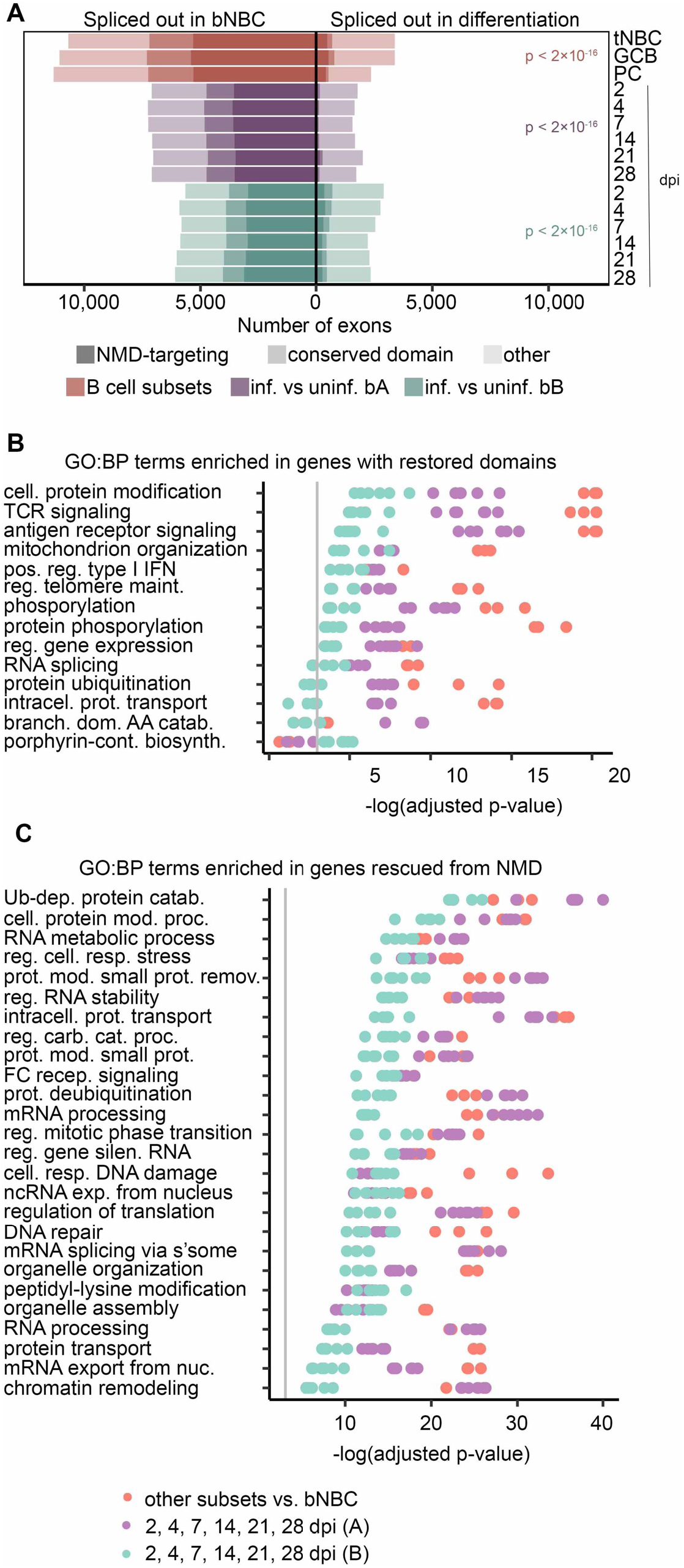
Exon inclusion restores domains and evades NMD in important pathways. (A) Exons with altered usage (rMATS FDR < 0.05) in lymphoid (tonsil) NBCs (tNBC), GC B cells (GCB) and PCs compared to blood NBCs (red) or EBV infection of blood B cells (purple = infection timecourse A, green = infection timecourse B). Exons that encode conserved protein domains are lightly shaded, exons that induce NMD targeting when skipped are heavily shaded. P-values for the proportion of damaging exon skipping events (i.e. NMD-targeting or domain-skipping) in blood NBCs compared to differentiating or EBV-infected cells were calculated using a pairwise comparison of proportions test with Bonferroni correction. (B) GO Biological Process term enrichment in genes with conserved protein domains restored by exon inclusion in differentiating or EBV-infected cells. The top 5 terms with lowest adjusted p-value for each experiment/timepoint are shown. (C) GO Biological Process term enrichment in genes with NMD targeting reduced by exon inclusion in differentiated or EBV-infected cells. The top 10 terms with lowest adjusted p-value for each experiment/timepoint are shown. For panels C-D, redundant terms are not displayed, complete enrichment information is available in Supplemental file 7. Vertical grey line indicates adjusted p-value = 0.05. Adjusted p-values calculated by Enrichr using Fisher’s exact test with Benjamini-Hochberg correction. For all panels B cell subset comparisons include 3 subsets (tonsil NBCs, GC B cells, and PCs) compared to blood NBCs. Each B cell subset includes 3 replicates. Blood infection timecourse A (purple) includes 6 timepoint comparisons with 3 replicates each, Blood infection timecourse B (green) includes 6 timepoint comparisons with 1 replicate each.

### Functional pathways impacted by restored transcripts in activated B cells

To predict cellular functions impacted by restoration of splicing fidelity, we separately assessed GO term enrichment in genes with altered mRNA splicing in each B cell subset compared to blood NBCs, and in EBV-infected cells at each infection timepoint compared to parental uninfected B cells. Results from these different experiments were remarkably consistent. Genes with transcripts whose restored exons encoded conserved domains were enriched for GO terms related to B cell activation and differentiation, such as *GO:0050851: antigen receptor-mediated signaling pathway* and *GO:0032481: positive regulation of type I interferon production;* and terms related to cellular signaling, such as *GO:0006468: protein phosphorylation*. Consistent with previous reports that splicing factors themselves are frequently alternatively spliced (34, 35), *GO:0008380: RNA splicing* and related terms were also enriched (**Figure 4B, Supplemental file 7**). Analysis of genes with transcripts rescued from NMD by exon inclusion in differentiating B cells revealed a set of enriched GO terms that partially overlapped those of genes with restored protein domains. Many of these genes are involved in *GO:0043470: regulation of carbohydrate catabolic process* and *GO:1901990: regulation of mitotic cell cycle phase transition*, suggesting a role for alternative splicing in supporting the cell growth and proliferation induced by both antigen-mediated activation and EBV infection. Splicing-related GO terms, such as *GO:0000398: mRNA splicing, via spliceosome*, were also enriched in genes with reduced mRNA NMD targeting (**Figure 4C, Supplemental file 7**). Genes with reduced mRNA intron retention in differentiating B cells, especially at the lymphoid NBC and GC B cell stages, were also enriched in terms related to mRNA splicing and processing (e.g. *GO:0000398: mRNA splicing, via spliceosome* and *GO:0006397: mRNA processing*; **Figure S5, Supplemental file 7**), consistent with previous findings in mice (7). The enrichment of activation-related GO terms in genes with altered mRNA splicing points to a multifaceted role for splicing fidelity enhancement in B cell differentiation. Furthermore, enrichment in splicing-related GO terms suggests a positive feedback mechanism whereby production of splicing factors themselves is improved, leading to further increases in splicing factor availability and function.

### Pre-activation splicing changes prepare naïve B cells for antigen encounter

Exon usage in blood B cells changes rapidly after EBV infection, with a majority of splicing alterations established by 2 dpi and very few further changes at later timepoints (**Figures 2, S2, 3A**). In contrast, mRNA abundance levels change dramatically both before and after 2 dpi (17, 18, 20), and major physiological changes occur later, in particular a phase of hyperproliferation that begins 3-4 dpi (21). Overall, fewer splicing changes occurred upon EBV infection of NBCs derived from lymphoid tissue than from blood.

While the Blueprint Epigenome samples represent snapshots of B cell subsets from different donors rather than a timecourse experiment, they can be arranged to reflect the progression of NBCs from blood to secondary lymphoid tissue, followed by differentiation into GC B cells and finally into PCs. Differentiation-associated splicing changes also occurred remarkably early, with blood NBCs displaying a singular splicing profile that was lost upon entry into secondary lymphoid tissue. Indeed, the lymphoid NBC splicing profile more closely resembled that of activated B cells than that of blood NBCs (**Figures 1A, 1D, 1E, S2A**). To investigate more closely the relationships of the B cell subsets we used rMATS to perform pairwise comparisons of blood NBCs, lymphoid NBCs and GC B cells to fully differentiated PCs. By far the largest number of exon usage differences was observed when comparing blood NBCs to PCs. Both lymphoid NBCs and GC B cells had relatively few exon usage differences compared to PCs (**Figure 5A**). The reduction in intron retention occurred more slowly, with a progressive stepwise reduction from blood NBCs to lymphoid NBCs and GC B cells relative to PCs (**Figure 5B**). Transcript abundance changes also occurred between blood and lymphoid NBCs, with the transition to a fully differentiated PC mRNA expression program that had begun but was by no means complete in lymphoid NBCs (**Figure 5C**). Because NBCs, unlike GC B cells and PCs, have not yet encountered their cognate antigen, this suggests that a majority of exon splicing changes occur upon NBC migration to secondary lymphoid tissue in preparation for -not in response to -antigen encounter and BCR signaling.

**Figure 5.**
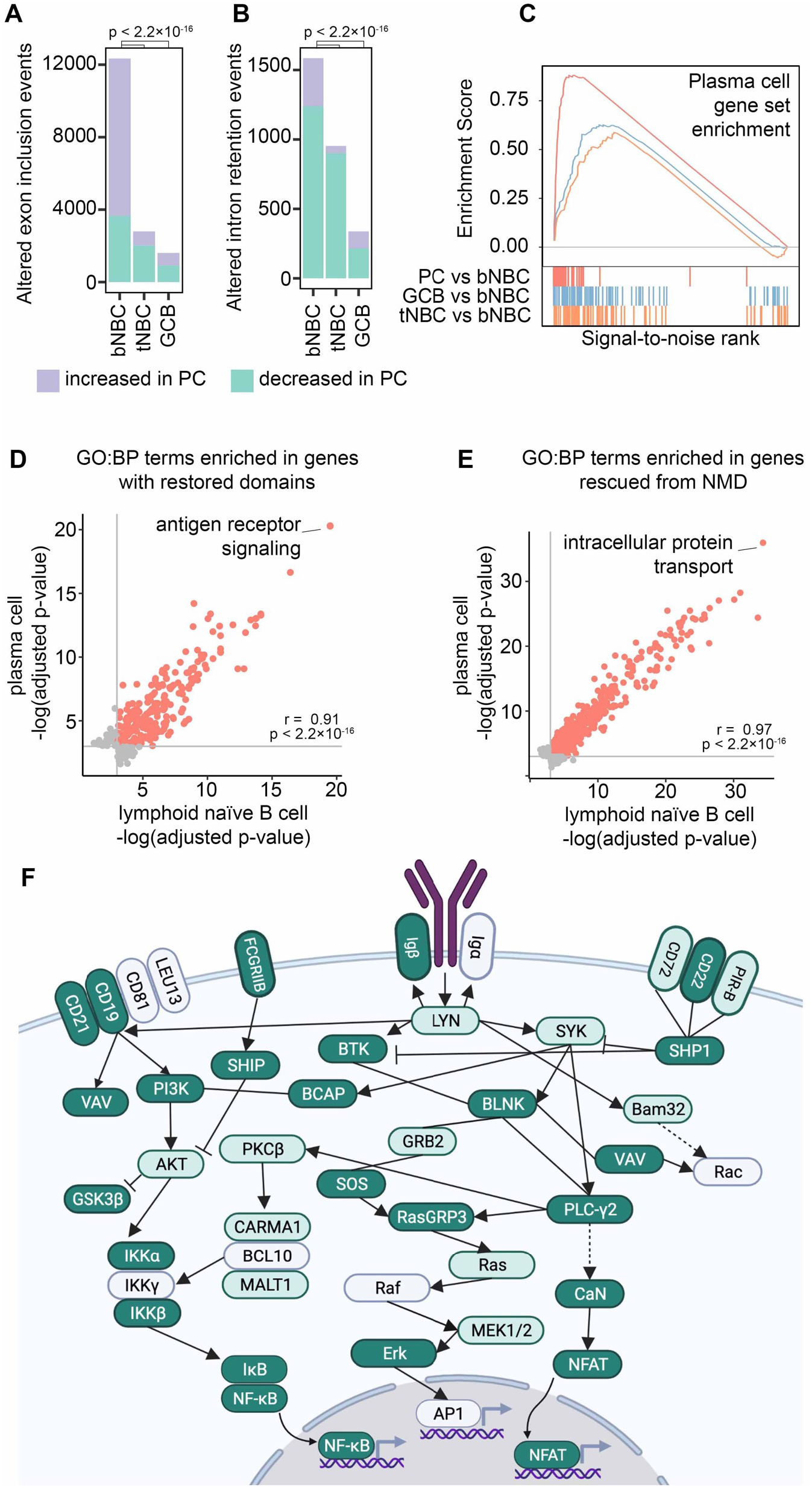
Lymphoid tissue-associated splicing repair prepares naïve B cells for antigen encounter. Exon usage changes (rMATS FDR < 0.05) in blood NBCs (bNBC), lymphoid (tonsil) NBCs (tNBC) and GC B cells (GCB) compared to PCs. P-values calculated by Pearson’s Chi-squared test with Yates’ continuity correction. (B) Intron retention changes (rMATS FDR < 0.05) in blood NBCs (bNBC), tonsil NBCs (tNBC) and GC B cells (GCB) compared to PCs. P-values calculated by Pearson’s Chi-squared test with Yates’ continuity correction. (C) Enrichment scores from GSEA of RNA abundance changes in PCs (red, maximum enrichment score = 0.880, p = 0), GC B cells (GCB, blue, maximum enrichment score = 0.627, p = 0.001) or tonsil NBCs (tNBC, orange, maximum enrichment score = 0.588, p = 0.021) compared to blood NBCs. Gene set: TARTE_PLASMA_CELL_VS_B_LYM PHOCYTE_UP from MSigDB. Colored vertical lines indicate genes in the gene set. (D) GO Biological Process term enrichment in genes with conserved protein domains restored by exon inclusion in tonsil NBCs or in PCs compared to blood NBCs. All significantly enriched terms (adjusted p-value < 0.05) are shown. Grey lines indicate adjusted p-value = 0.05. Red points: significant enrichment in both PCs and tonsil NBCs. Grey points: significant enrichment in tonsil NBCs or PCs only. Correlation (r) and p-value calculated using Pearson’s product-moment correlation. Full GO term information is shown in Supplemental file 8..(E) GO Biological Process term enrichment in genes with NMD targeting reduced by exon inclusion in tonsil NBCs or in PCs compared to blood NBCs. All significantly enriched terms (adjusted p-value < 0.05) are shown. Grey lines indicate adjusted p-value = 0.05. Red points: significant enrichment in both PCs and tonsil NBCs. Grey points: significant enrichment in tonsil NBCs or PCs only. Correlation (r) and p-value calculated using Pearson’s product-moment correlation. Full GO term information is shown in Supplemental file 8. (F) BCR signaling pathway (KEGG). Genes in dark green show altered exon usage (rMATS FDR <0.05) in tonsil NBCs compared to blood NBCs and in EBV-infected cells compared to parental blood NBCs (both experiments, all timepoints). Genes in light green show altered exon usage in tonsil NBCs compared to blood NBCs and in at least one EBV infection experiment timepoint. Image created with Biorender.com. Full information is shown in Supplemental file 9.

To further assess the similarity of lymphoid NBC and PC splicing profiles we examined GO term enrichment in genes with altered mRNA splicing relative to blood NBCs. GO term enrichment in these comparisons was strikingly similar, both when examining genes with protein domains restored and genes with mRNA NMD targeting reduced by B cell (pre-)activation (**Figure 5D-E**). One of the GO terms with the strongest enrichment in genes with restored protein domains was *GO:0050851: antigen receptor-mediated signaling pathway* (**Supplemental file 8**), further supporting a potential role for splicing fidelity enhancement in preparing cells for antigen encounter and BCR signaling. An in-depth examination of mRNA splicing changes in the BCR signaling pathway (36) revealed that an extraordinary number of genes in this pathway showed a change in exon usage as NBCs entered the secondary lymphoid tissue (**Figure 5F** and **Supplemental file 9**). Few further exon usage changes occurred upon differentiation into GC B cells or PCs (**Supplemental file 9**), suggesting that changes at the mRNA splicing level contribute to preparing the BCR signaling pathway for activation at the protein level.

## Discussion

We used RNA-seq datasets from four different human B cell subtypes, each with multiple donors, to examine stage-specific splicing profiles during B cell differentiation. Blood-derived NBCs had a remarkable number of unannotated splicing events with 2127 high-confidence novel splice junctions detected consistently across all donors and over 30% of expressed genes containing at least one novel splicing event (**Figure 1A**). While statistically significant splicing changes were observed at each stage of B cell differentiation, the preponderance of changes occurred in the blood NBC to lymphoid NBC transition. Assessing the functional significance of alternative splicing in blood NBCs, we found that a high percentage of exon skipping events specific to blood NBCs are predicted to undergo nonsense mediated RNA decay or to exclude a known conserved domain (**Figure 4A**), suggesting that the transition to the lymphoid microenvironment increases splicing fidelity in preparation for activation. Using EBV infection as a direct model of B cell activation, in blood B cells we similarly found that exon skipping and intron retention were substantially reduced (i.e. splicing fidelity increased) prior to B cell proliferation, and in lymphoid NBCs the remaining lower-level splicing infidelity was further corrected, again prior to EBV-induced proliferation (**Figure 2**).

The striking changes in exon usage between blood NBCs and lymphoid NBCs, with exon usage in lymphoid NBCs more closely resembling that in antigen-experienced, fully differentiated PCs, were an unexpected finding. These changes precede the major physiological changes of activation and are accompanied by the earliest changes in gene expression levels (**Figure 5C**, (37, 38)) The previous observation that, unlike the other B cell subsets, the two sets of NBCs do not significantly differ in DNA methylation (38) suggests that the lymphoid NBC samples in the Blueprint Epigenome project are unlikely to be significantly contaminated by activated B cells and do in fact represent a distinct population of antigen-naïve cells. B cell “pre-activation” at the RNA level is supported by our observations that adenoid NBCs show fewer splicing changes upon EBV infection than blood B cells, despite the fact that the blood B cells used in both EBV infection timecourses were not explicitly enriched for naïve cells, although they likely consisted largely of NBCs (39). Further support comes from the observations of others that EBV infection-induced proliferation proceeds much more quickly in adenoid B cells than in blood B cells (40). These parallel observations of NBCs from tonsils and from adenoids suggest that exon usage changes are a common feature of NBCs entering secondary lymphoid tissues throughout the body, though splicing changes associated with lymph nodes, spleen or gut lymphoid tissues remain to be resolved.

The progression from blood NBC to GC B cell and PC constitutes a major coordinated overhaul of the cell, from DNA methylation (38) and chromatin structure (41) through gene expression (42) and up to the physiological level. A global increase in splicing efficiency appears to be well-integrated into this complex process, with a decrease in the transcriptomic “noise” of aberrantly-spliced, nonfunctional transcripts accompanying the overall increase in transcription. This is likely supported by a suite of biological processes, potentially including increased transcription and improved exon/intron usage in splicing factors, optimized RNA polymerase II transcription rates, improved snRNA and snoRNA biogenesis mediated by AQR or other factors, and/or increased NMD efficiency as reported in mouse activated B cells (43).

The specific cellular, or in the case of infection, viral factors that trigger the widespread differentiation-induced increases in splicing fidelity remain unclear, but likely involve a coordinated array of signaling events. The pervasive similarity of splicing changes between *in vitro* EBV infection and *in vivo* B cell differentiation suggests a common mechanism, with the virus hijacking a cellular system rather than directly intervening in the splicing process. Importantly, splicing changes occur early in infection and are complete by 2 dpi, before most viral genes reach their full expression level (17, 18, 20). Among the viral genes with high early expression is Epstein Barr virus nuclear antigen 2 (EBNA2). EBNA2 interacts with cellular splicing factors and has the capacity to impact splicing (44). However, infection of blood B cells with recombinant EBV lacking EBNA2 induced a large-scale shift in cellular exon splicing very similar to that induced by wild-type EBV (**Figure S6**), despite very different impacts on host mRNA levels (20) and a complete abrogation of infection-associated proliferation and immortalization (21). Similarly, the viral EBNA-LP gene is expressed early after infection and has in fact been demonstrated to control a subset of cellular splicing changes, but the majority of infection-associated splicing changes are not dependent on EBNA-LP expression (45). Interestingly, EBNA2 and EBNA-LP expression, together with physical binding of the viral protein gp350 to the cellular complement receptor CR2 (also known as CD21) have been shown to be sufficient to drive proliferation in blood NBCs (46). While the splicing profile of these gp350/EBNA2/EBNA-LP-stimulated cells is not known, it raises the possibility that infection-induced splicing changes are triggered not by viral gene expression but by the virion’s physical engagement of CR2 and the resultant cellular signaling cascade.

CR2 signaling is also known to enhance antigen-mediated B cell activation (47). While the secondary lymphoid tissues are rich in antigens and immune cells, greatly increasing the chances that an NBC will encounter both its antigen and a T cell with affinity for the same antigen, these tissues are also enriched in complement proteins. Although BCR affinity for antigens is highly specific and the chances of an NBC encountering its antigen at any given time are relatively low, CR2 has generic affinity for the abundant complement protein fragment C3dg, suggesting that NBCs may encounter and bind complement proteins before or in the absence of their cognate antigens. CR2 is also stimulated by CD23 (48) and IFN-α (49), both present in the secondary lymphoid tissues (50). While it is not yet known whether complement signaling impacts splicing, this pathway represents an intriguing candidate for the initiation of the pre-proliferative splicing fidelity enhancement that is common to both B cell differentiation and EBV infection.

## Methods

### RNA-Seq data acquisition and analysis

RNA-Seq datasets from human donor B cell subsets were obtained from the European Genome-Phenome Archive (EGA) with permission from the Blueprint Epigenome Consortium. RNA-Seq datasets from GM12878 and infection of primary and immortalized B cells were obtained from the European Nucleotide Archive (ENA). RBP-knockdown datasets were obtained from ENCODE. Accession numbers for all datasets are in Supplemental file 1. Data from donor 2 from blood B cell infection B were found to be corrupted and were not used. Data from one donor in the lymphoid NBC infection at 8 dpi were reported as not passing quality controls; this dataset was excluded from both the original analysis (17) and the present one.

### RNA-Seq datasets were analyzed as follows

1. Reads were aligned and mapped with STAR v2.5.2a (51) with the following settings, optimized for splicing analysis: --chimSegmentMin 2 –outFIlterMismatchNmax 3 –alignEndsType EndToEnd – outSAMstrandField intronMotif –alignSJDBOverhangMin 6 – alignIntronMax 300000
2. For analysis of exon inclusion level and statistical assessment of splicing changes (skipped exon, retained intron, alternate 3’ splice site, alternate 5’ splice site and mutually exclusive exons) between conditions, the BAM alignment files produced by STAR were analyzed with rMATS 4.1.1 or v3.1.0 (52).
3. For analysis of intron retention level in each sample, the BAM alignment files produced by STAR were analyzed with IRFinder v1.3.0 (53).
4. Gene expression levels were calculated as Transcripts per Million (TPM) using kallisto v0.46.0 (54).
5. Differential gene expression was calculated using sleuth (55).
6. Gene Set Enrichment Analysis (GSEA) was performed in GSEA 3.0 (56) using a signal-to-noise rank and the TARTE_PLASMA_CELL_VS_B_LYMPHOCYTE_UP gene set.

Alignment, mapping and splicing analysis used a reference genome containing the GRCh38 assembly of the human genome and the EBV genome (Akata strain, NCBI accession number KC207813.1, (57)) and a reference transcriptome annotation containing the Ensembl 90 human transcript annotation and EBV (Akata) transcript annotation (58).

### Identification and analysis of unannotated splice junctions

Canonical splice junctions (with a GT-AG splicing pattern) identified by STAR were assigned to genes in the Ensembl 90 human transcriptome annotation using the findOverlaps function in GRanges (59). For each robustly expressed (TPM > 10) gene, the annotated splice junction with the highest number of uniquely mapping reads was identified. Then, unannotated splice junctions with uniquely mapping reads equaling at least 10% that of that gene’s highest-depth annotated splice junction were selected for further analysis (Supplemental file 2).

Donor and acceptor chromosome coordinates of unannotated splice junctions from each replicate were compared to each other in a pairwise fashion to calculate the proportion of shared unannotated splice junctions (the number of common unannotated splice junctions divided by the total number of unannotated splice junctions in the two datasets being compared). A hypergeometric test was used to evaluate the statistical significance of the shared set of unannotated junctions.

### Evaluation of exon commitment and intron retention

To determine the exon commitment level for each sample, the exon inclusion level (“IncLevel”) of each exon in the rMATS output from genes with TPM > 10 was extracted. In the case of duplicate exons (i.e., the same exon being skipped by multiple alternate splice junctions), the minimum reported inclusion level was used. The exon commitment score for the sample was calculated as:

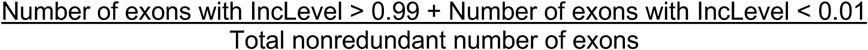

To determine the global intron retention level, in each sample the “IRratio” was extracted for each intron in IRFinder output for genes with TPM > 10. The number of genes with at least one intron with IRratio > 0.25 was then divided by the total number of genes with TPM > 10.

### Cross-dataset splicing alteration comparisons

For pairwise comparisons of splicing profile changes across experiments (B cell differentiation, EBV infection and ENCODE RBP knockdowns), all chromosome coordinates of high-confidence significantly altered splice junctions (FDR < 0.0005) in rMATS output from each comparison were compared to each other in a pairwise fashion to calculate the proportion of splice junctions altered in both conditions. A hypergeometric test was used to evaluate the statistical significance of the shared set of altered junctions.

### Splice site strength, intron and exon length, GC content

The sets of exons with significantly higher inclusion (rMATS FDR 0.05, IncLevelDifference > 0) and introns with lower retention (rMATS FDR 0.05, IncLevelDifference < 0) in PCs or EBV infected cells compared to uninfected blood NBCs from the corresponding dataset were examined. Splice site strength was calculated using MaxEntScan (26).

### Identification of NMD-targeting exon-skipping events

Alternately spliced exons (rMATS FDR 0.05) that had nucleotide lengths not divisible by 3 and were in annotated coding sequences were analyzed for NMD-targeting potential. Reading frames for each possible affected isoform of the affected gene were scanned for premature stop codons (PTCs). If a PTC was detected >50 nt upstream of a splice junction, the exon-skipping event was determined to be potentially NMD-targeting.

### Examination of conserved protein domain (CD) changes

Alternately spliced exons (rMATS FDR 0.05) that had nucleotide lengths divisible by 3 and were in annotated coding sequences were analyzed for CD changes. The encoded amino acid sequences of the exons were obtained using a standard codon table and the amino acid sequences were submitted to the NCBI CD-Search Tool (Conserved Domain database, Expect Value threshold 0.01 with composition-corrected scoring in automatic search mode).

### Functional enrichment analysis

To examine genes with unannotated splicing in B cell subsets, genes with identified unannotated splicing (see subsection “*Identification and analysis of unannotated splice junctions*”) were compared across replicates. Genes that contained unannotated splice junctions in all replicates of a subset were used for functional enrichment analysis. To examine genes with differentiation-associated alternative splicing, lists of genes with CD restoration, NMD targeting or intron retention were identified as described as above.

Lists of genes were analyzed with EnrichR (15). GO Biological Process 2021 results were downloaded and plots created with R/ggplot2. GO terms with Adjusted p-value < 0.05 were considered to be significantly enriched.

## Acknowledgements

This study makes use of data generated by the Blueprint Consortium. A full list of the investigators who contributed to the generation of the data is available from www.blueprint-epigenome.eu. Funding for the project was provided by the European Union’s Seventh Framework Programme (FP7/2007-2013) under grant agreement no 282510 – BLUEPRINT. This study also makes use of data generated by the ENCODE consortium, in particular by the Brenton Graveley laboratory at the University of Connecticut.

This research was supported in part using high performance computing (HPC) resources and services provided by Technology Services at Tulane University, New Orleans, LA. We also acknowledge the use of computational resources and expertise from the Tulane Cancer Crusaders Next Generation Sequence Analysis Core at the Tulane Cancer Center.

This work was supported by the NIH grants R01CA262090, R01CA243793, and P01CA214091 (EKF). NAU was supported by a fellowship from the Leukemia & Lymphoma Society.

## References

1. J. M. Rodriguez, F. Pozo, T. di Domenico, J. Vazquez, M. L. Tress, An analysis of tissue-specific alternative splicing at the protein level. PLoS Comput Biol 16, e1008287 (2020).

2. E. Furlanis, P. Scheiffele, Regulation of Neuronal Differentiation, Function, and Plasticity by Alternative Splicing. Annu Rev Cell Dev Biol 34, 451–469 (2018).

3. L. Yue, R. Wan, S. Luan, W. Zeng, T. H. Cheung, Dek Modulates Global Intron Retention during Muscle Stem Cells Quiescence Exit. Dev Cell 53, 661–676 e666 (2020).

4. C. R. Edwards et al., A dynamic intron retention program in the mammalian megakaryocyte and erythrocyte lineages. Blood 127, e24–e34 (2016).

5. H. Pimentel et al., A dynamic intron retention program enriched in RNA processing genes regulates gene expression during terminal erythropoiesis. Nucleic Acids Res 44, 838–851 (2016).

6. L. Chen et al., Transcriptional diversity during lineage commitment of human blood progenitors. Science 345, 1251033 (2014).

7. S. Ullrich, R. Guigo, Dynamic changes in intron retention are tightly associated with regulation of splicing factors and proliferative activity during B-cell development. Nucleic Acids Res 48, 1327–1340 (2020).

8. A. A. Pai et al., Widespread Shortening of 3’ Untranslated Regions and Increased Exon Inclusion Are Evolutionarily Conserved Features of Innate Immune Responses to Infection. PLoS Genet 12, e1006338 (2016).

9. T. Ni et al., Global intron retention mediated gene regulation during CD4+ T cell activation. Nucleic Acids Res 44, 6817–6829 (2016).

10. J. J. Wong et al., Orchestrated intron retention regulates normal granulocyte differentiation. Cell 154, 583–595 (2013).

11. L. Chen et al., Genetic Drivers of Epigenetic and Transcriptional Variation in Human Immune Cells. Cell 167, 1398–1414 e1324 (2016).

12. D. Adams et al., BLUEPRINT to decode the epigenetic signature written in blood. Nat Biotechnol 30, 224–226 (2012).

13. H. Cho et al., High-resolution transcriptome analysis with long-read RNA sequencing. PLoS One 9, e108095 (2014).

14. H. Tilgner, F. Grubert, D. Sharon, M. P. Snyder, Defining a personal, allele-specific, and single-molecule long-read transcriptome. Proc Natl Acad Sci U S A 111, 9869–9874 (2014).

15. Z. Xie et al., Gene Set Knowledge Discovery with Enrichr. Curr Protoc 1, e90 (2021).

16. D. A. Thorley-Lawson, EBV Persistence--Introducing the Virus. Curr Top Microbiol Immunol 390, 151–209 (2015).

17. P. Mrozek-Gorska et al., Epstein-Barr virus reprograms human B lymphocytes immediately in the prelatent phase of infection. Proc Natl Acad Sci U S A 116, 16046–16055 (2019).

18. C. Wang et al., RNA Sequencing Analyses of Gene Expression during Epstein-Barr Virus Infection of Primary B Lymphocytes. J Virol 93 (2019).

19. T. Inagaki et al., Direct Evidence of Abortive Lytic Infection-Mediated Establishment of Epstein-Barr Virus Latency During B-Cell Infection. Front Microbiol 11, 575255 (2020).

20. Y. Yanagi et al., RNAseq analysis identifies involvement of EBNA2 in PD-L1 induction during Epstein-Barr virus infection of primary B cells. Virology 557, 44–54 (2021).

21. D. Pich et al., First Days in the Life of Naive Human B Lymphocytes Infected with Epstein-Barr Virus. mBio 10 (2019).

22. E. L. Van Nostrand et al., A large-scale binding and functional map of human RNA-binding proteins. Nature 583, 711–719 (2020).

23. E. P. Consortium, An integrated encyclopedia of DNA elements in the human genome. Nature 489, 57–74 (2012).

24. C. A. Davis et al., The Encyclopedia of DNA elements (ENCODE): data portal update. Nucleic Acids Res 46, D794–D801 (2018).

25. F. Kouzine et al., Global regulation of promoter melting in naive lymphocytes. Cell 153, 988–999 (2013).

26. G. Yeo, C. B. Burge, Maximum entropy modeling of short sequence motifs with applications to RNA splicing signals. J Comput Biol 11, 377–394 (2004).

27. N. Fong et al., Pre-mRNA splicing is facilitated by an optimal RNA polymerase II elongation rate. Genes Dev 28, 2663–2676 (2014).

28. C. E. Zorca et al., Myosin VI regulates gene pairing and transcriptional pause release in T cells. Proc Natl Acad Sci U S A 112, E1587–1593 (2015).

29. U. Braunschweig et al., Widespread intron retention in mammals functionally tunes transcriptomes. Genome Res 24, 1774–1786 (2014).

30. P. A. Galante, N. J. Sakabe, N. Kirschbaum-Slager, S. J. de Souza, Detection and evaluation of intron retention events in the human transcriptome. RNA 10, 757–765 (2004).

31. N. J. Sakabe, S. J. de Souza, Sequence features responsible for intron retention in human. BMC Genomics 8, 59 (2007).

32. E. Nagy, L. E. Maquat, A rule for termination-codon position within intron-containing genes: when nonsense affects RNA abundance. Trends Biochem Sci 23, 198–199 (1998).

33. S. Lu et al., CDD/SPARCLE: the conserved domain database in 2020. Nucleic Acids Res 48, D265–D268 (2020).

34. L. F. Lareau, S. E. Brenner, Regulation of splicing factors by alternative splicing and NMD is conserved between kingdoms yet evolutionarily flexible. Mol Biol Evol 32, 1072–1079 (2015).

35. N. J. Homa et al., Epstein-Barr virus induces global changes in cellular mRNA isoform usage that are important for the maintenance of latency. J Virol 87, 12291–12301 (2013).

36. M. Kanehisa, S. Goto, KEGG: kyoto encyclopedia of genes and genomes. Nucleic Acids Res 28, 27–30 (2000).

37. K. Chokeshai-u-saha, C. Lepoivre, L. Grieco, C. Nguyen, K. Ruxrungtham, Comparison of immunological characteristics of peripheral, splenic and tonsilar naive B cells by differential gene expression meta-analyses. Asian Pac J Allergy Immunol 30, 326–330 (2012).

38. M. Kulis et al., Whole-genome fingerprint of the DNA methylome during human B cell differentiation. Nat Genet 47, 746–756 (2015).

39. H. Morbach, E. M. Eichhorn, J. G. Liese, H. J. Girschick, Reference values for B cell subpopulations from infancy to adulthood. Clin Exp Immunol 162, 271–279 (2010).

40. R. Zeidler, P. Meissner, G. Eissner, S. Lazis, W. Hammerschmidt, Rapid proliferation of B cells from adenoids in response to Epstein-Barr virus infection. Cancer Res 56, 5610–5614 (1996).

41. K. R. Kieffer-Kwon et al., Myc Regulates Chromatin Decompaction and Nuclear Architecture during B Cell Activation. Mol Cell 67, 566–578 e510 (2017).

42. A. Tesi et al., An early Myc-dependent transcriptional program orchestrates cell growth during B-cell activation. EMBO Rep 20, e47987 (2019).

43. A. Tinguely et al., Cross talk between immunoglobulin heavy-chain transcription and RNA surveillance during B cell development. Mol Cell Biol 32, 107–117 (2012).

44. Q. Peng et al., Epstein-Barr virus EBNA2 phase separation regulates cancer-associated alternative RNA splicing patterns. Clin Transl Med 11, e504 (2021).

45. E. Manet et al., Modulation of alternative splicing during early infection of human primary B lymphocytes with Epstein-Barr virus (EBV): a novel function for the viral EBNA-LP protein. Nucleic Acids Res 49, 10657–10676 (2021).

46. A. J. Sinclair, I. Palmero, G. Peters, P. J. Farrell, EBNA-2 and EBNA-LP cooperate to cause G0 to G1 transition during immortalization of resting human B lymphocytes by Epstein-Barr virus. EMBO J 13, 3321–3328 (1994).

47. R. A. Barrington et al., CD21/CD19 coreceptor signaling promotes B cell survival during primary immune responses. J Immunol 175, 2859–2867 (2005).

48. J. P. Aubry, S. Pochon, P. Graber, K. U. Jansen, J. Y. Bonnefoy, CD21 is a ligand for CD23 and regulates IgE production. Nature 358, 505–507 (1992).

49. A. X. Delcayre et al., Epstein Barr virus/complement C3d receptor is an interferon alpha receptor. EMBO J 10, 919–926 (1991).

50. K. Sandberg, M. L. Eloranta, I. L. Campbell, Expression of alpha/beta interferons (IFN-alpha/beta) and their relationship to IFN-alpha/beta-induced genes in lymphocytic choriomeningitis. J Virol 68, 7358–7366 (1994).

51. A. Dobin et al., STAR: ultrafast universal RNA-seq aligner. Bioinformatics 29, 15–21 (2013).

52. S. Shen et al., rMATS: robust and flexible detection of differential alternative splicing from replicate RNA-Seq data. Proc Natl Acad Sci U S A 111, E5593–5601 (2014).

53. R. Middleton et al., IRFinder: assessing the impact of intron retention on mammalian gene expression. Genome Biol 18, 51 (2017).

54. N. L. Bray, H. Pimentel, P. Melsted, L. Pachter, Near-optimal probabilistic RNA-seq quantification. Nat Biotechnol 34, 525–527 (2016).

55. H. Pimentel, N. L. Bray, S. Puente, P. Melsted, L. Pachter, Differential analysis of RNA-seq incorporating quantification uncertainty. Nat Methods 14, 687–690 (2017).

56. A. Subramanian et al., Gene set enrichment analysis: a knowledge-based approach for interpreting genome-wide expression profiles. Proc Natl Acad Sci U S A 102, 15545–15550 (2005).

57. Z. Lin et al., Whole-genome sequencing of the Akata and Mutu Epstein-Barr virus strains. J Virol 87, 1172–1182 (2013).

58. T. O’Grady et al., Global transcript structure resolution of high gene density genomes through multi-platform data integration. Nucleic Acids Res 44, e145 (2016).

59. M. Lawrence et al., Software for computing and annotating genomic ranges. PLoS Comput Biol 9, e1003118 (2013).

