## Supplemental figures for "Reversal of splicing infidelity is a pre-activation step in B cell differentiation"

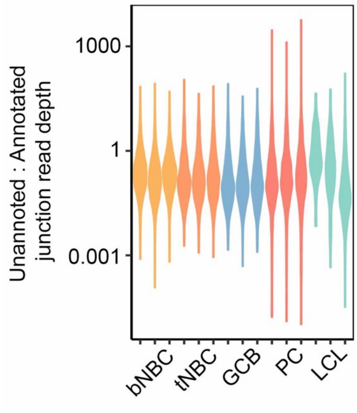


**Figure S1. Unannnotated splicing occurs at a range of depths compared to annotated splicing**. Ratios of the number of reads uniquely aligning to unannotated splice junctions to the number of reads uniquely aligning to annotated splice junctions that share their donors or acceptors. In the case of more than one annotated splice junction sharing donor/acceptor sites with the unannotated junction, the annotated splice junction with the highest read depth is used. bNBC = blood naïve B cells, tNBC = tonsil naïve B cells, GCB = germinal center B cells, PC = plasma cells, LCL = lymphoblastoid cell line GM12878.

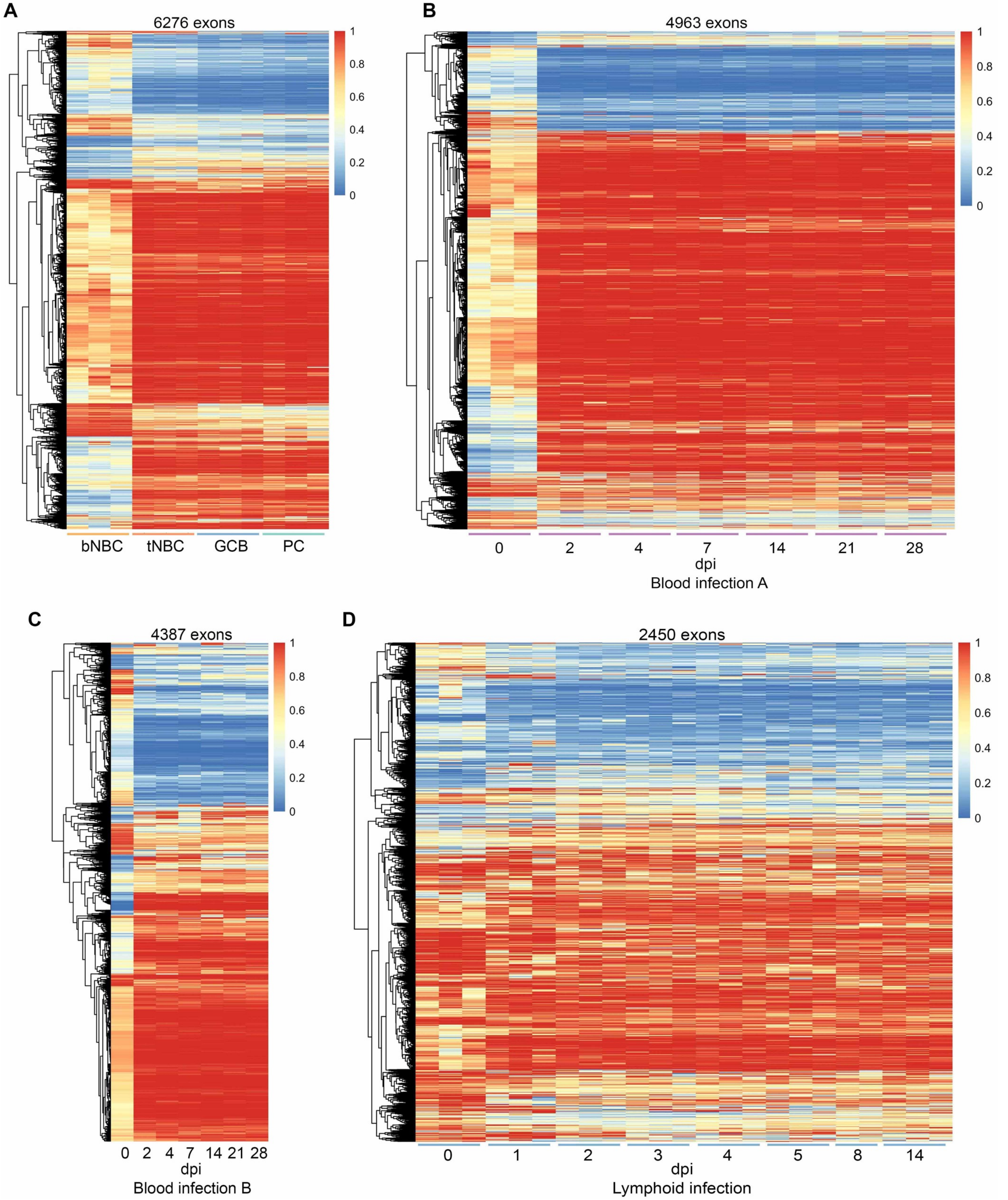


**Figure S2. Exon usage changes occur early in differentiation/infection and do not revert.** (A) Exon inclusion level (rMATS IncLevel) for exons in genes with TPM > 10 showing a high-confidence usage change (rMATS FDR < 0.0005, |IncLevelDifference| ≥ 0.2) in lymphoid (tonsil) naïve blood cells (tNBC), GC B cells (GCB) or plasma cells (PC) compared to blood naïve B cells (bNBC). N = 3 donors per subset.

*Continued next page*

*Continued from previous page:*

**Figure S2. Exon usage changes occur early in differentiation/infection and do not revert …** (B-C) Exon inclusion level (rMATS IncLevel) for exons in genes with TPM > 10 showing a high-confidence usage change (rMATS FDR < 0.0005, |IncLevelDifference| ≥ 0.2) in any timepoint compared to 0 dpi for (B) EBV infection of blood B cells timecourse A, N = 3 replicates, (C) EBV infection of blood B cells timecourse B, N = 1 replicate, and (D) EBV infection of lymphoid (adenoid) NBCs, N = 3 replicates.

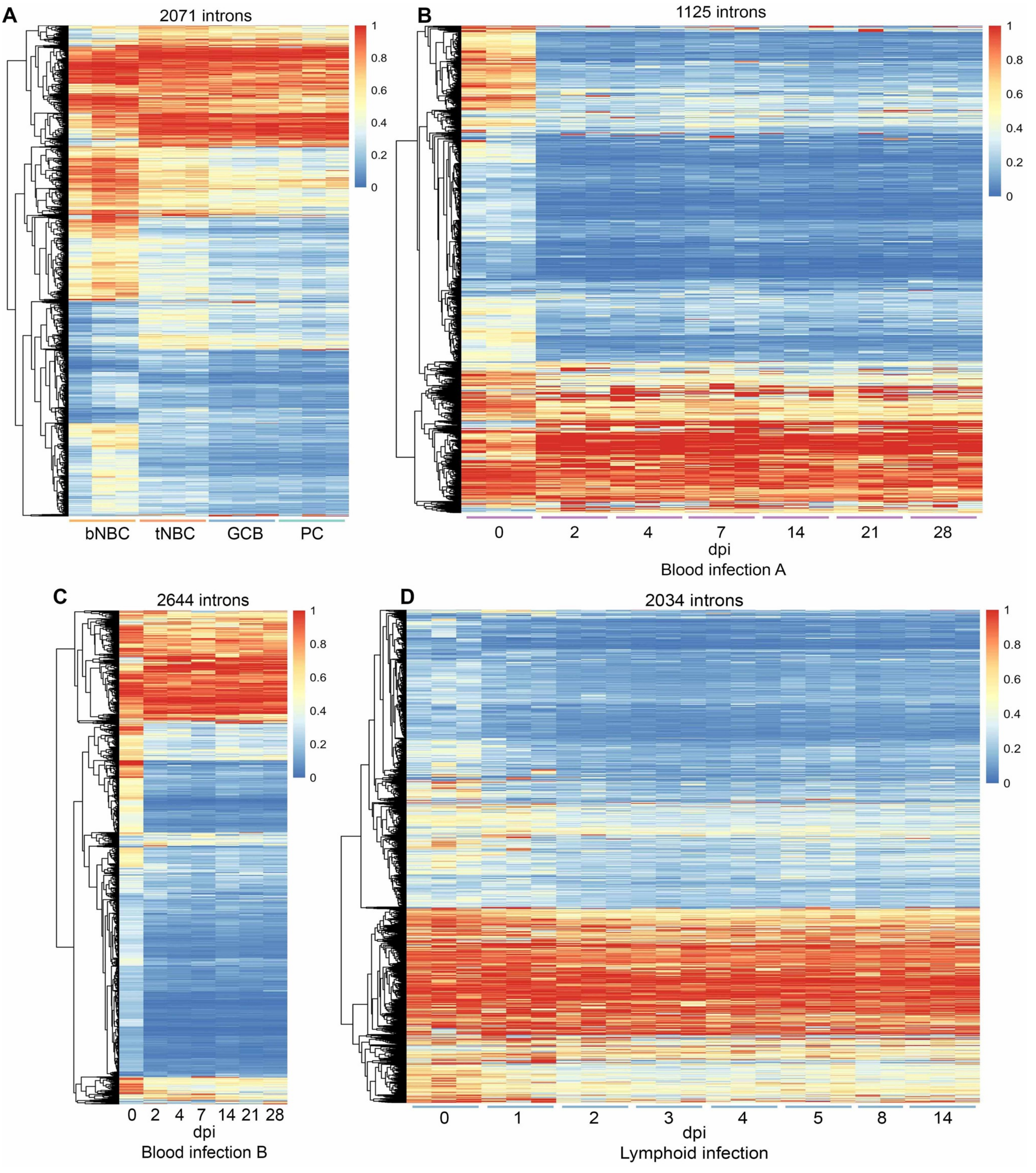


**Figure S3: Intron usage changes in differentiation/infection** (A) Intron retention level (rMATS IncLevel) for introns in genes with TPM > 10 showing a change in retention (rMATS FDR < 0.05, |IncLevelDifference| ≥ 0.1) in lymphoid (tonsil) NBCs (tNBC), GC B cells (GCB) or plasma cells (PC) compared to blood NBCs (bNBC). N = 3 donors per subset. (B-D) Intron retention level (rMATS IncLevel) for introns in genes with TPM > 10 showing a change in retention (rMATS FDR < 0.05, |IncLevelDifference| ≥ 0.1) in any timepoint compared to 0 dpi for (B) EBV infection of blood B cells timecourse A, N = 3 replicates, (C) EBV infection of blood B cells timecourse B, N = 1 replicate, and (D) EBV infection of lymphoid (adenoid) NBCs, N = 3 replicates. Hierarchical clustering was performed and heatmaps generated using pheatmap.

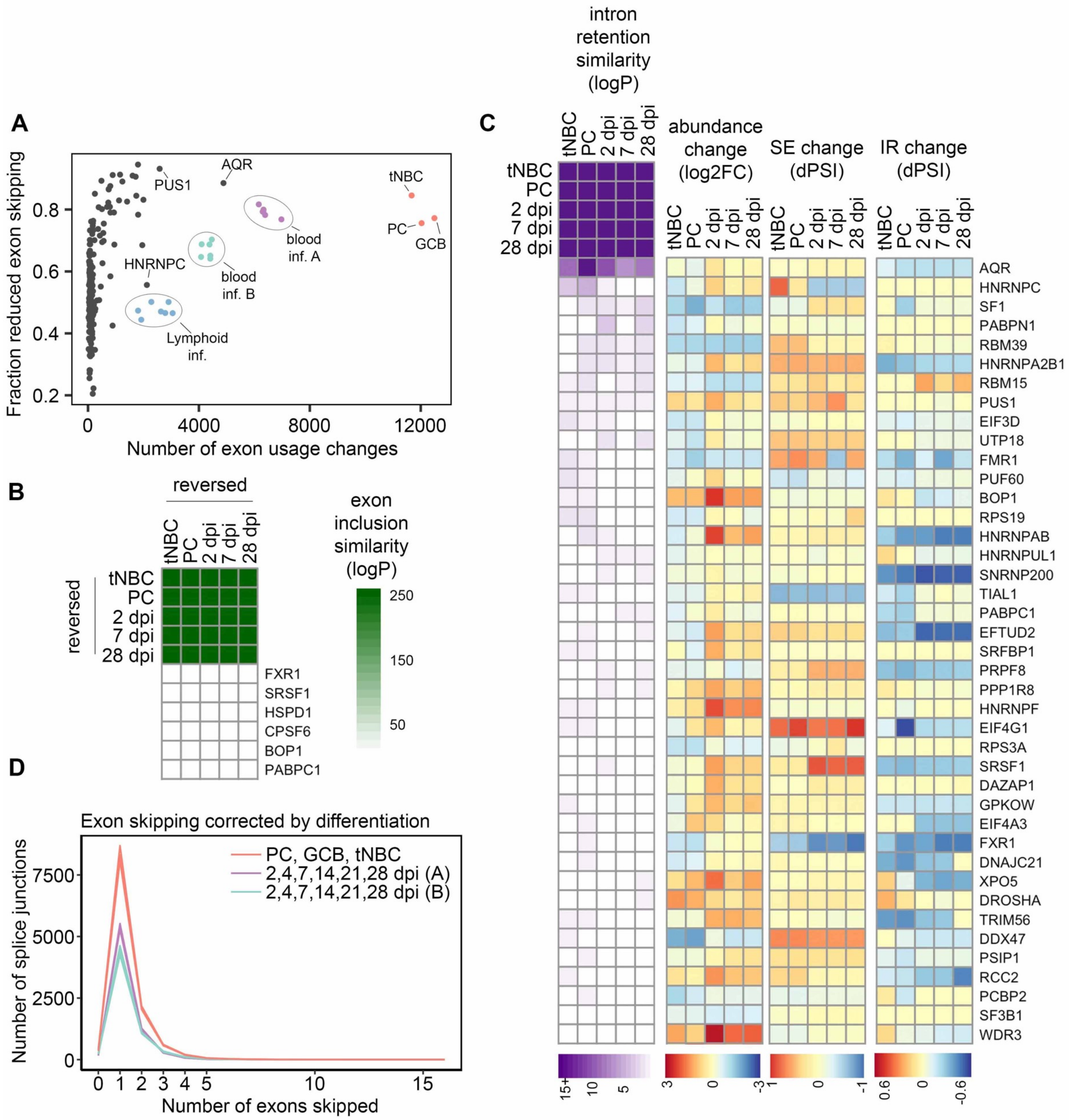


**Figure S4. Increased splicing program efficiency alters the splicing profile.** (A) Exon usage changes in differentiating B cells compared to blood NBCs (3 replicates per subset, ~$100\times10^6$ reads per replicate), EBV-infected cells compared to uninfected cells (Blood A: 6 timepoints, 3 replicates per timepoint, 13-$85\times10^6$ reads per replicate; Blood B: 6 timepoints, 1 replicate per timepoint, ~$60\times10^6$ reads per replicate; Lymphoid: 7 timepoints, 3 replicates per timepoint, 120-$165\times10^6$ reads per replicate), and siRNA knockdowns of 182 RBPs compared to control K562 cells (2 replicates per RBP, 25-$30\times10^6$ reads per replicate). (B) Heatmap indicating exon inclusion similarity for RBPs with exon inclusion changes most similar to the artificially reversed values for B cell differentiation/EBV infection. (C) Heatmaps indicating intron retention decrease similarity (logP = -log(p-value) by hypergeometric test), abundance change (log2FC = $\log_2$(fold change) by sleuth), skipped exon (SE) change (dPSI = difference in percent spliced-in by rMATS), and intron retention (IR) change (dPSI by IRFinder) for RBPs with intron retention changes most similar to those of B cell differentiation/EBV infection. For RBP genes with multiple splice junctions the SE and IR events with the highest absolute dPSI value measurable in all compared replicates are displayed. PC = PCs compared to blood NBCs; tNBC = tonsil NBCs compared to blood NBCs; 2, 7 and 28 dpi = EBV-infected cells at specified timepoints compared to uninfected cells in Blood timecourse A.

*Continued next page*

*Continued from previous page:*

**Figure S4. Increased splicing program efficiency alters the splicing profile.** … (D) Number of exons skipped by exon-skipping splice junctions that are corrected by differentiation or EBV infection (rMATS FDR $< 0.05$). B cell subset comparisons include 3 subsets (tonsil NBCs, GC B cells, and PCs) compared to blood NBCs. Each B cell subset includes 3 replicates. Blood infection timecourse A (purple) includes 6 timepoint comparisons to 0 dpi with 3 replicates each, Blood infection timecourse B (green) includes 6 timepoint comparisons to 0 dpi with 1 replicate each.

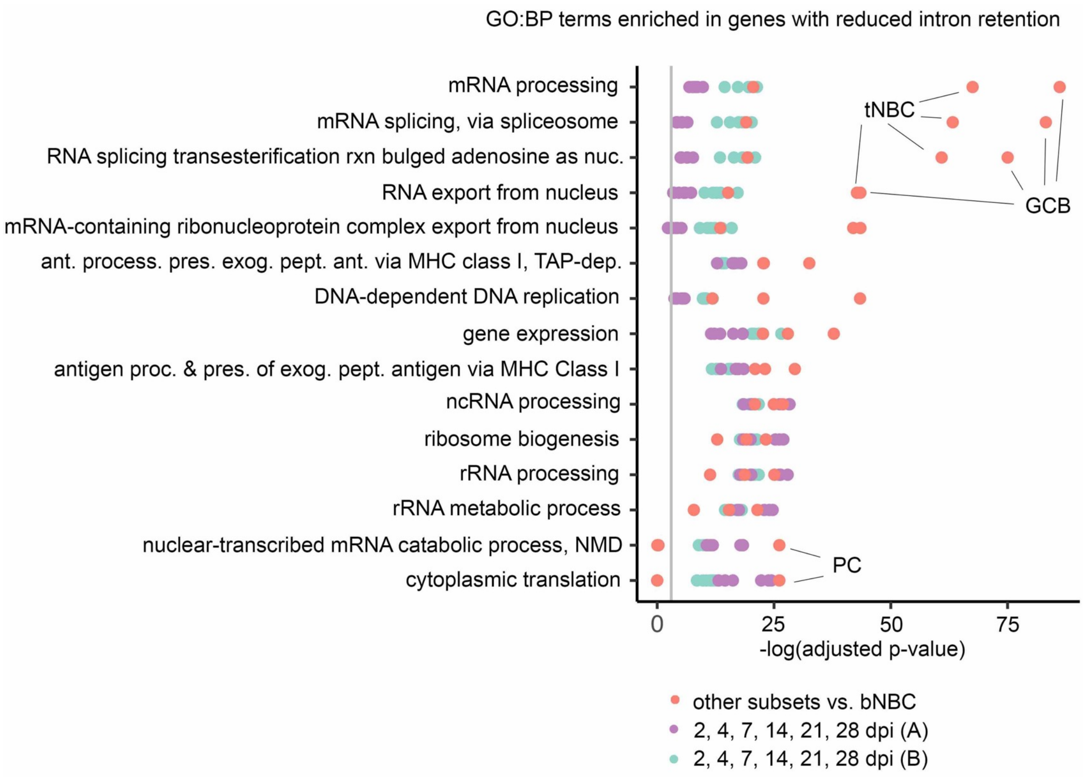


**Figure S5. Functions of genes with a loss of intron retention.** GO Biological Process term enrichment in genes with reduced intron retention in differentiating or EBV-infected cells. Top 5 terms with the lowest adjusted p-value for each experiment/timepoint are shown. Complete enrichment information available in Supplemental file 7. Vertical grey line indicates adjusted p-value = 0.05. Adjusted p-values calculated by Enrichr using Fisher's exact test with Benjamini-Hochberg correction.

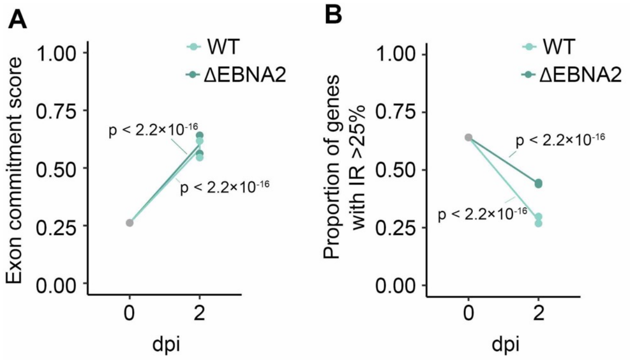


**Figure S6. EBV EBNA2 has minimal effects on exon usage changes.** (A) Exon commitment score for blood NBCs infected with wild-type (WT) EBV or EBV with deleted EBNA2 gene (ΔEBNA2). Exon commitment score = proportion of exons either fully included (rMATS IncLevel > 0.99) or fully excluded (rMATS IncLevel < 0.01) in genes with TPM > 10. P-values calculated relative to uninfected cells using 2-sample tests for equality of proportions with continuity correction. 0 dpi = 1 replicate, 2 dpi = 2 replicates. (B) Proportion of genes containing introns with high retention levels (IRFinder IRratio > 0.25) for blood NBCs infected with wild-type (WT) EBV or EBV with deleted EBNA2 gene (ΔEBNA2). P-values calculated using 2-sample tests for equality of proportions with continuity correction. 0 dpi = 1 replicate, 2 dpi = 2 replicates.
